# Arsenic induced alteration of neuroregulatory gene expression in *Macrobrachium lamarrei*: a neurotoxicological approach to grooming behaviour

**DOI:** 10.1101/2022.08.30.505863

**Authors:** Chayan Munshi, Alpana Mukhuty, Arindam Bandyopadhyay, Arpan Dey Bhowmik, Paritosh Mondal, Pallab Shaw, Shelley Bhattacharya

**Author notes:** Corresponding author: Chayan Munshi (Email address).

## Abstract

Arsenic is a highly active neuromodulator which can induce neurological disorders in invertebrates. The present study is a neurotoxicological approach to assess the toxicity of arsenic trioxide, where grooming behaviour is considered as a behavioural biomarker of arsenic induced stress in the prawns. Non-lethal exposure to arsenic trioxide, manipulates the expression pattern of neuroregulatory marker genes in a freshwater prawn, *Macrobrachium lamarrei* and induces symptoms of autism spectrum disorders (ASDs) in a short time.

## Introduction

Animal behaviour is the manifestation of neuronal activities in an organism where analysis of behavioural phenotypes is considered as a reliable tool to understand neuronal functions. Grooming is a neurophenotypic stereotyped behaviour, established as a behavioural index of stress and neurological disorders in crustacean and mammals **(Munshi and Bhattacharya, 2020; Kalueff et al., 2016)**. The present study reveals a neurogenetic alteration caused by arsenic trioxide in *Macrobrachium lamarrei*, a freshwater prawn, which stimulates repetitive grooming behaviour (possible indication of autism spectrum disorder) in the species. *Macrobrachium lamarrei* (Arthropoda: Decapoda: Crustacea) is a reliable model to study behavioural toxicity which is a sensitive aquatic invertebrate species known for evident acute and robust behavioural response in different experimental conditions. Its abundance in all types of freshwater bodies and response to diverse environmental conditions presents it as a promising aquatic bioindicator model. The aim of the study was to analyse the effect of a well-reported neurotoxicant and environmental pollutant, arsenic trioxide, in altering normal expression level of marker neuroregulatory genes which control the neuronal activities thus manifesting behavioural patterns in the organism. We have considered grooming as the behavioural index and hypothesized possible alterations in neurogenetic expressions associated with it. Accordingly, three experimental groups were set up: a) prawns collected from natural habitat, b) prawns reared in uncontaminated water in the laboratory (control) and c) prawns exposed to arsenic (As) contamination in the laboratory. To the best of our knowledge, this is the first-ever study that attempts to unravel the underlying neuro-regulation of enhanced grooming activity in prawns as a function of non-lethal arsenic exposure.

Arsenic contamination in water has developed into a serious threat to aquatic organisms. Even non-lethal concentration of arsenic can acutely or chronically influence the normal physiology and behaviour of animals owing to the incessant environmental exposure to this chemical. Neurologists are emphasising on behavioural analysis and considering behavioural indices or markers to assess stress induced neurological disorders. **Munshi et al., 2021** have demonstrated that non-lethal dose of arsenic trioxide can trigger repetitive grooming behaviour in the prawns within 24 hours of exposure. An ethogram study of this organism (supplementary data) showed that grooming is the most robust behaviour which is altered (enhanced) notably due to arsenic contamination. Our study focuses on neurexin-neuroligin and acetylcholinesterase genes in this organism. Moreover, the stress has been assessed by heat shock protein genes (stress markers).

## Experimental design and methodology

Prawns were collected from native ponds, which are normally used for fishing. Ten prawns were sacrificed for gene expression assessment. Rest, a population of twenty prawns were collected and reared in clean filtered tap water in the laboratory, maintaining a consistent temperature of 25 ± 2°C (approximate pond temperature). Food (flour globule) was provided *ad libitum*. After an acclimatization period of 20 days, ten random prawns from the same population were transferred to the arsenic contaminated water. Arsenic trioxide was provided at a concentration of 1.7 mg L^−1^ **(Munshi and Bhattacharya, 2020)**. After 24 hours of exposure, the prawns were sacrificed for the gene expression assessment.

TRIzol® TRI Reagent (Sigma) was used to extract total RNA according to the manufacturer’s instructions. Tissue samples (dissected brain) were homogenised in TRI Reagent (1 ml per 50–100 mg of tissue). Chloroform was added and the samples were shaken vigorously for 15 seconds, allowed to stand and then centrifuged at 12,000 × g for 15 minutes at 2-8 °C. The colourless upper aqueous phase containing RNA was transferred to a fresh tube and 2-propanol was added. The mixtures were centrifuged at 12,000 × g for 10 minutes at 2-8 °C. The RNA precipitate was washed once with 75% ethanol, then centrifuged at 12,000 × g for 5 minutes and dried at room temperature. The RNA pellet was re-suspended in nuclease-free water (20 μL). RNA pellet was treated with DNase I (Invitrogen) to remove genomic DNA if any. RNA quantity and purity were evaluated using a spectrophotometer (Nanodrop2000c, Thermo Fisher Scientific) at 260 nm (200 ng RNA/sample). Complementary DNA (cDNA) was synthesized using a Revert Aid first strand cDNA synthesis kit (Fermentas) with 1 μg of RNA. The cDNAs were adjusted to 100 μL with sterile water. For Real-time PCR, each reaction was prepared with a cDNA sample (50 ng), 10 pM of gene-specific primers, SYBR green master mix (Invitrogen) and nuclease free water to a final volume of 10 μL. For the first PCR cycle, the reaction mixture was initially denatured at 95°C for 10 min (1 cycle), followed denaturing for 15 s at 95°C, annealing at 60°C for 30 s, and extension at 72°C (30 s) for a total of 40 cycles. Fluorescence from SYBR stacked DNA was measured during annealing step. Dissociation step (95°C for 15s, 60°C for 30s, 95°C for 15s) was conducted for validation of amplification of a single product in each PCR reaction. Gapdh (glyceraldehyde-3-phosphate dehydrogenase) was used as a reference gene to normalize the target genes. To obtain the relative quantities of genes, threshold cycle (ct) values were used. Data were analysed using the 2-ΔΔct method. In the negative control, no cDNA template was used. Significant differences among gene expression levels in various treatments were analysed using Quant Studio 5 software. One way analysis of variance was used to compare the mean values.

## Results and discussion

Heat Shock Proteins (HSPs) are highly conserved and present in all living organisms, but they vary in their expression pattern. The induction of HSPs occurs under various environmental stress situations including temperature, heavy metals, bacterial and viral infections, nutrient deprivation, hydrostatic pressure, pollutants, UV exposure, oxygen radicals, malignant agents, and hypoxia. Due to stimulation in different stressful situations, HSPs are collectively known as stress proteins. The principal HSPs are grouped into conserved classes based on their molecular weight and sequence homology. These include hsp110, hsp100, hsp90, hsp70, hsp60, hsp40 and some smaller molecular weight HSPs **(Feder and Hofmann, 1999)**. The widespread expression of HSPs in insects indicates their significant role in insect adaptability to a fluctuating environment. Hsp60, 70 and 90 have a significant role as molecular chaperones in subcellular organelles and stabilise proteins to regulate correct folding by aggregation of proteins into an oligomeric configuration. These proteins are also involved in many cellular processes including signal transduction, DNA replication, protein synthesis and protein trafficking **(Otaka et al., 1994)**. Greater knowledge of the changes in expression and transcription patterns of HSPs and related proteins could provide a clearer insight into the responses of prawns to stress at cellular and physiological levels.

Our study confirmed the differential expression of *hsp10, hsp60, hsp70* and *hsp90* in the prawn brain from the natural habitat, control and arsenic contaminated groups. *HSPs* are stress markers. Expressions of *hsp10* and *hsp90* are downregulated in the prawns from both natural habitat and arsenic contaminated water in respect of the control. In natural habitat, the reduction of *hsp10* and *hsp90* is 4.1 and 2.8 folds respectively (significant, p<0.05); in arsenic contaminated water, the reduction is 5.5 and 3.8 folds respectively (significant, p<0.05). Brain *hsp60* and *hsp70* are upregulated in the prawns from both natural habitat and arsenic contaminated water, in respect to the control. In natural habitat, the increase in the expression levels of *hsp60* and *hsp70* is 7.4 folds and 18.3 folds respectively (significant, p<0.05); in arsenic contaminated water, the increment is 7.4 folds and 14.0 folds respectively (significant, p<0.05). The level of *hsp60* in natural habitat and arsenic contaminated water is similar but for *hsp70*, the level in prawns from natural habitat is 1.3 fold more than those in arsenic contaminated water **(Figure 1)**. Therefore, in the prawns from both natural habitat and arsenic contaminated water, have significant expression level of *HSPs*.

**Figure 1.**
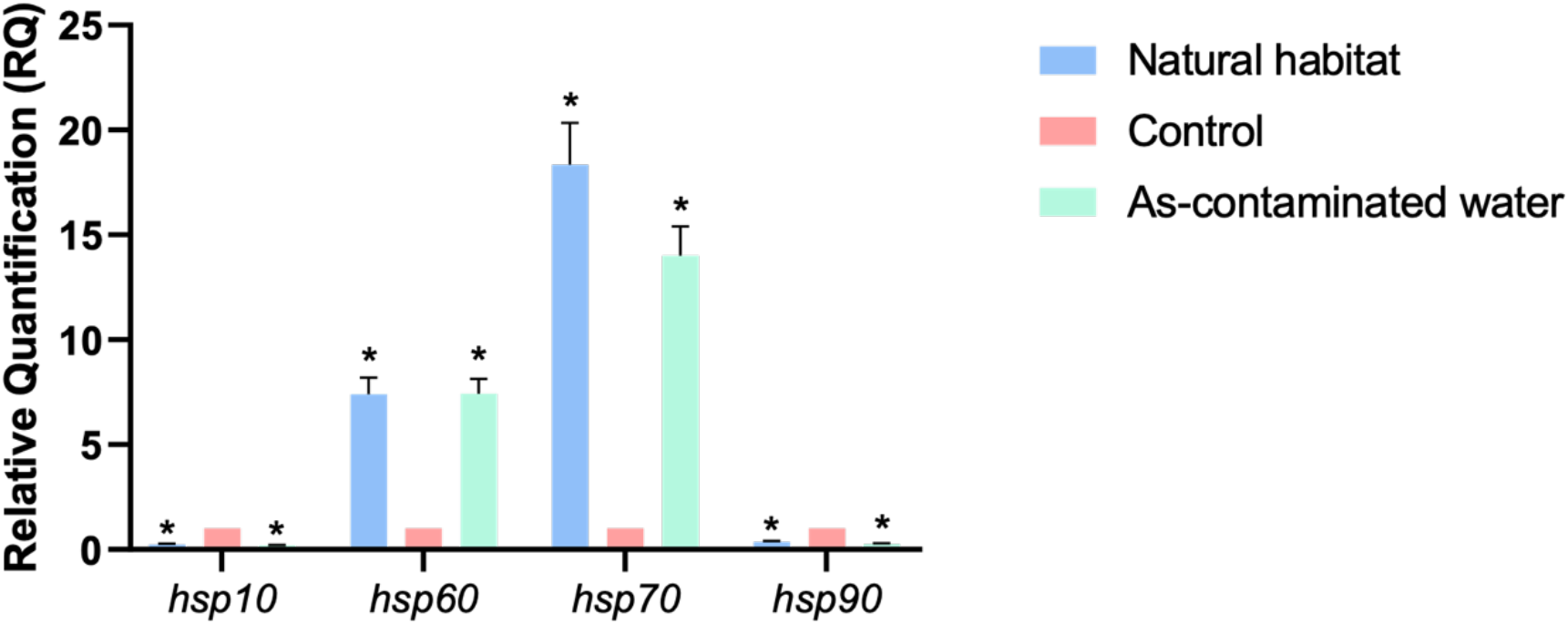
Expression levels of *hsp10, hsp60, hsp70* and *hsp90* in the brain of *Macrobrachium lamarrei*, reared in natural habitat and the laboratory (control and arsenic contaminated water). Values are presented as the mean±S.D. Significance level (p<0.05) is indicated as * in the figure.

Neurexin-neuroligin adhesion complex in the trans-synaptic region of neurons plays a crucial role in neurodevelopment, synapse specification, maintenance and sensory regulation. Neurexins and neuroligins are studied extensively in the area of neuronal disorders. Structurally, neuroligins resemble acetylcholinesterase (AChE), which is an essential neuro-enzyme, known to execute a critical role in neurotransmission **(Biswas et al., 2008)**. Normally, single neurexin is found in invertebrates which binds to diverse ligands. On the other hand, there are four types of neuroligins in vertebrates and five types in invertebrates **(Sudhof, 2017; Biswas et al., 2008)**. Sudhof (2017) summarized in a review that neuroligin-1 is concentrated in excitatory synapses, neuroligin-2 in inhibitory, dopaminergic, cholinergic synapses, neuroligin-3 in both excitatory and inhibitory synapses and neuroligin-4 in glycinergic synapses.

Acetylcholinesterase (AChE) is one of the most crucial enzymes regulating neurotransmission and is present in the synapse of cholinergic neurons in both vertebrates and invertebrates. There are two types of AChE present in the brain of invertebrates, AChE-1 and AChE-2. From the perspective of evolution, AChE-2 is a possible substitute for AChE-1 **(Kim and Lee, 2013)**. AChE activity is considered to evaluate neurotoxicity in animals. AChE-1 is a regulator of biotic and abiotic stress in honeybee; where stress has been found to affect the expression level of AChE-1 **(Kim et al., 2019)**. Expression level of two types of acetylcholinesterase genes (*ache*): *ache1* and *ache2* were analysed. Both the types of these enzymes are downregulated in natural habitat and arsenic contaminated water, compared to the control. The decrease in the expression of *ache1* is 3.8 folds in the natural habitat and 5.8 folds in arsenic contaminated water (significant, p<0.05). The decrease in the expression of *ache2* is 2.0 folds in natural habitat and 4.3 folds in arsenic contaminated water (significant, p<0.05). However, in both the conditions, expression of *ache2* is higher than that of *ache1* **(Figure 2)**.

**Figure 2.**
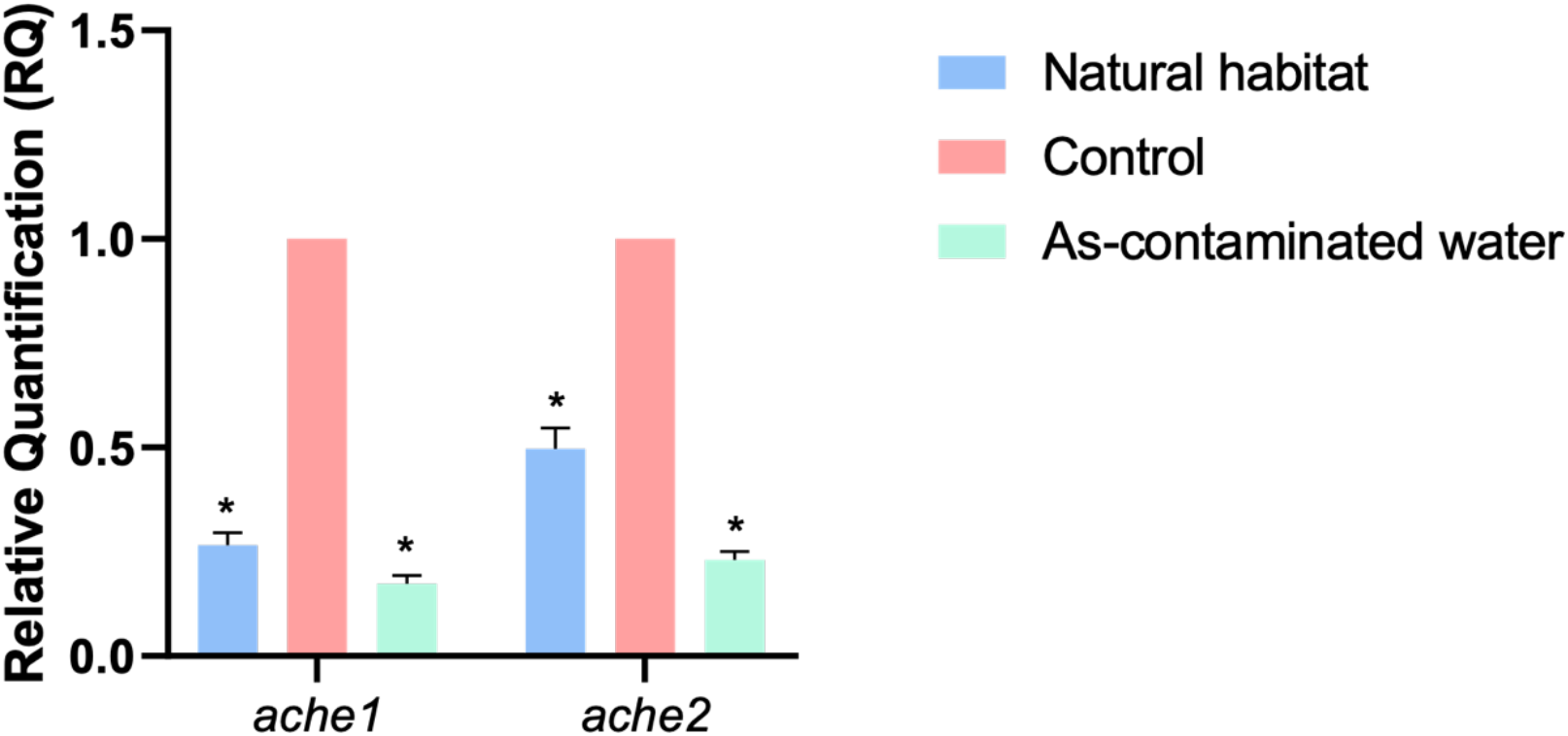
Expression levels of *ache1* and *ache2* in the brain of *Macrobrachium lamarrei*, reared in natural habitat and the laboratory (control and arsenic contaminated water). Values are presented as the mean±S.D. Significance level (p<0.05) is indicated as * in the figure.

The expression level of neurexin1 gene (*nrx1*) in the prawns from both natural habitat and arsenic contaminated water is upregulated, against the control. The expression significantly (p<0.05) increased by 2.2 folds in prawns from natural habitat and 5.6 folds in arsenic contaminated water. Besides, analysis of the expression and function of neuroligins is a way to understand synapse-activities. These proteins are used as a marker of neuropsychiatric or neurological disorders.

Neuroligin-1 gene (*nlg1*) is downregulated in the group of prawns from both natural habitat and arsenic contaminated water, compared to the control. The decrease in the expression level is 1.9 folds in natural habitat and 4.7 folds in arsenic contaminated water (significant, p<0.05). Deletion of *nlg1* can cause extreme reiteration of behaviour (like grooming) which is a prospective Autism Spectrum Disorder **(Blundell et al., 2010)**; similarly reduction in the expression of this gene caused a momentous enhancement in grooming activity (within 24 hours), which was reported earlier by **Munshi et al., 20201**. Deficiency of *nlg1* in *Caenorhabditis elegans* showed hypersensitivity to toxic elements like mercury **(Hunter et al., 2010)**.

Neuroligin-2 gene (*nlg2*) is upregulated in the group of prawns from both natural habitat and arsenic contaminated water, compared to the control. The increase in the expression level is only 0.6 fold in natural habitat but 4.4 folds (significant, p<0.05) in the arsenic contaminated water. Deletion of *nlg2* in mice can lead to anxiety like behaviour, caused by decrease in inhibitory synaptic function **(Blundell et al., 2009)**. However, overexpression of *nlg2* in mice can cause increased synaptic contact size, anxiety and impaired social communication **(Hines et al., 2008)**. This gene is also reported to be a key player in neuro-stress susceptibility **(Heshmati et al., 2017)**.

Neuroligin-3 gene (*nlg3*) and neuroligin-5 gene (*nlg5*) are upregulated in the natural habitat, but downregulated in the arsenic contaminated water, in comparison to the control. In natural habitat, the enhancements in the expression level of *nlg3* and *nlg5* are 2.4 and 4.5 folds (significant, p<0.05); in arsenic contaminated water, the decrease in *nlg3* is 1.3 folds and in *nlg5* is 2.9 folds (significant, p<0.05). Deletion of *nlg3* can cause repetitive behaviour in mice **(Rothwell et al., 2014)**. In invertebrates (*Drosophila* as the study organism) *nlg1, nlg2* and *nlg3* are crucial for synapse maintenance and maturation. Downregulation of *nlg2* and *nlg4* can result in impairment of social behaviour in *Drosophila melanogaster* **(Corthal et al., 2017)**. Deficiency in *nlg1, nlg2* and *nlg3* can result in severe impairment in synaptic transmission **(Sudhof, 2017)**. In sensory depleted condition, honeybee shows alterations in patterns of neurexin-neuroligins **(Biswas et al., 2010)**.

Neuroligin-4 gene (*nlg4*) is upregulated in the group of prawns from both natural habitat and arsenic contaminated water against the control; the expression level in natural habitat is 2.0 fold higher (significant, p<0.05). However, the upregulation in arsenic contaminated water is not notable. Deletion of *nlg4* results in reduced social interactions in mice **(Jamain et al., 2008)**.

Hence, the analysis of different genes considered in this study aided in understanding the arsenic mediated neurogenetic regulation in the brain of prawns.

Therefore, the expression of neuroligins in pond water is in the following order:

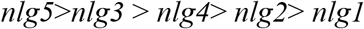

and expression of neuroligins in arsenic contaminated water is in the following order:

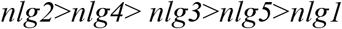

To summarize, the patterns of neurexin-neuroligin expression data in prawns from natural habitat show upregulation of *nrx1* and all *nlgs* except *nlg1*. In arsenic contaminated water there is a drastic increase in *nrx1* and *nlg2* **(Figure 3)**.

**Figure 3.**
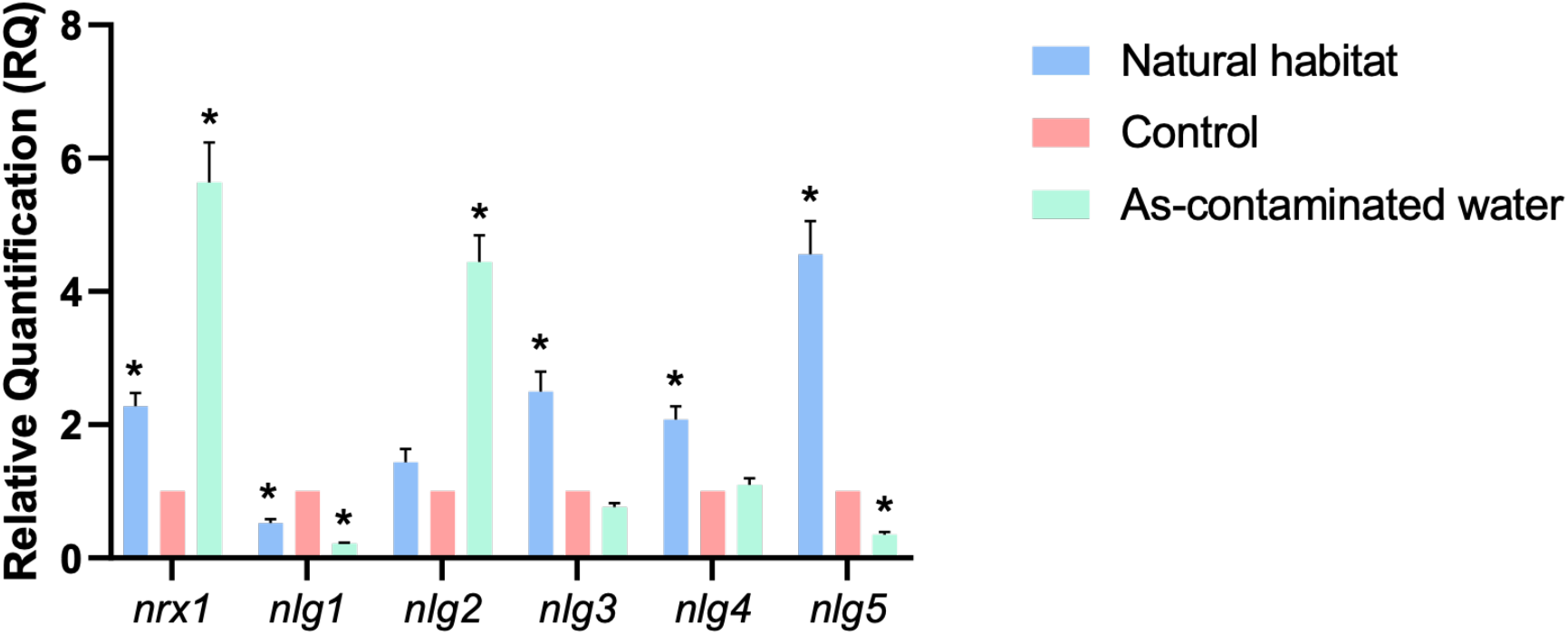
Expression levels of *nrx1, nlg1, nlg2, nlg3, nlg4, nlg5* in the brain of *Macrobrachium lamarrei*, reared in natural habitat and the laboratory (control and arsenic contaminated water). Values are presented as the mean±S.D. Significance level (p<0.05) is indicated as * in the figure.

## Conclusion

The present data make it abundantly clear that arsenic is the key stressor to modulate neuro-regulatory mechanism governing several behavioural activities, which include notable grooming alterations, in *M. lamarrei*. In natural habitat, expressions of *hsp60* and *hsp70* are significantly higher than those in the control which indicates stress response in the organism. The pond water is free from pollutants (analysed earlier), hence it is presumed that chemical stressors do not induce stress in the prawns, rather there may be several biotic factors that stimulate HSP expressions. However, prawns in arsenic contaminated water (laboratory condition) demonstrate high rates of *hsp60* and *hsp70* expression within 24 hours, which is, however, less than or equal to that of the expression level in prawns from natural habitat. Moreover, our findings indicate that arsenic stress, in a short time (24 hours) can robustly alter the neuro-regulatory gene expression in the prawn brain. It can downregulate the genes such as *ache1, ache2, nlg1, nlg3* and *nlg5*. Previous reports on *nlg1* and *nlg3* deficiency in mice have shown repetitive behaviour and socio-behavioural impairments respectively, both of which are indices of autism spectrum disorders (ASDs). **Etherton et al., (2009)** showed that *nrx1* deficiency can induce repetitive grooming in mice. On the contrary, in the present investigation, we found overexpression of *nrx1* enhanced grooming activity, as found in honey bee **(Hamiduzzaman et al., 2017)**. The downregulation of *nlg1* strongly validates repetitive grooming activity. As *M. lamarrei* is not a social animal, deficiency of *nlg3* did not indicate any social impairment. Apart from grooming, other robust behaviours were not notably altered in this exposure time. Unlike mice, despite of the overexpression of *nlg2*, no aggression was observed in this organism. It is concluded that in arsenic contaminated state there will be only *nlg2* (overexpressed) to adhere to *nrx1* (overexpressed) in *M. lamarrei*. We are confirming that this model could elucidate a new area of biotic and abiotic stress related neuro-genetic regulation and neurological disorders in aquatic invertebrates.

## Acknowledgement

The authors gratefully acknowledge National Academy of Sciences, India for providing research grant in the NASI Honorary Scientist Scheme (Grant Number: NAS/96/5/2016-2017).

